# Towards a Physiologically Rigorous Heartbeat Dynamics Foundation Model

**DOI:** 10.64898/2026.11.24.768987

**Authors:** Gayathri Sunil, Bhuvana Ravi Kumar, Bharath Ramsundar, Sandya Subramanian

## Abstract

Capturing long-range heartbeat dynamics accurately from wearable electrocardiogram (ECG) recordings is a key challenge in personalized healthcare and autonomic neuroscience. Although classical methods are physiologically grounded and statistically rigorous, they cannot learn across population data and therefore cannot scale. Time series foundation models, on the other hand, can scale, but fail to capture physiologically meaningful beat-to-beat dynamics. We present, to our knowledge, the first probabilistic foundation model of heartbeat dynamics: a density-based neural temporal point process (NTPP) pretrained on RR-interval sequences from 11,226 subjects across five public datasets spanning normal sinus rhythm, arrhythmia, atrial fibrillation, and ischemia. We establish a physiological scaling law in this domain, showing that held-out metrics improve predictably with cohort size. We additionally show that scaling behavior in stochastic output domains like physiology should be evaluated by distributional calibration metrics (KS distance, goodness-of-fit) rather than point-error (MAE, RMSE), because point errors approach their irreducible floor while distributional properties continue to improve. Our pretrained model transfers zero-shot to a held-out ambulatory dataset, reaching 94.7% goodness-of-fit with no subject-specific data. In 5-minute autoregressive simulation, the model remains calibrated (87–88% goodness-of-fit) and generates trajectories with real physiological variability. Finally, we show that the learned representations are physiologically interpretable. The recurrent history embeddings separate normal and arrhythmia windows in held-out test subjects with a balanced accuracy of 95.7%. The lognormal mixture distributions show that arrhythmia subjects differ mainly in predictive spread (3.6x larger standard deviation than normal cohort), with only a small shift toward shorter mean RR than the normal group (0.79 vs. 0.86 s). These results establish population-scale pretraining as a foundation for physiologically grounded modeling and simulation of heartbeat dynamics.

## I. Introduction

We need computational models that can bridge the gap between modern artificial intelligence and long-established physiological knowledge. With such models, we could learn trends across large populations, generate novel clinical hypotheses across heterogeneous cohorts, and lay the foundation for patient-specific digital twins and in silico clinical trials. Human heartbeats are a good example of such a process, where the mechanisms that govern heartbeat generation have been extensively studied, and there are a large number of ECG datasets that contain valuable information about the health and functioning of both the cardiovascular and nervous systems. Human heartbeats are not deterministic; they are the result of a probabilistic process that generates events roughly every second. The temporal dynamics of this sequence are known to inform the underlying physiological state of an individual.

Human heartbeats can be described in a physiologically faithful way as a temporal point process: each beat is a discrete probabilistic event driven by the heart’s inherent pacemaker and complex physiological feedback loops. Despite having a mathematically rigorous framework to describe heartbeats, developing a robust generative computational model for heartbeat dynamics remains an open challenge, due to various overlapping physiological processes that govern heartbeat dynamics. Existing approaches [1], [2] that use classical, physiologically grounded point-process frameworks provide rigorous statistical characterization of heartbeat dynamics, but are fit independently to individual subjects, and, as is, don’t support learning a shared representation across subjects. They therefore cannot leverage the available large, heterogeneous ECG datasets available to learn a population model. On the other hand, ECG foundation models [3], [4] have demonstrated the potential of learning from large, heterogeneous cohorts, but focus primarily on waveform or morphology. While ECG waveforms provide useful information about the electrical activity of the heart, the temporal dynamics that inform autonomic regulation cannot be directly modeled using such approaches. Another modeling approach is that of time series foundation models [5], [6], which have shown strong performance for forecasting and learning generalizable temporal patterns across domains, but these fail to capture the physiology-specific long-range temporal structure of heart beats, including inherent beat-to-beat variability [7]. As these models are not designed with a physiological prior, they struggle to accurately characterize the stochastic nature of the heartbeat generation process.

Therefore, we need a unified modeling approach that can

(1) learn from large patient cohorts spanning diverse disease states, (2) generalize to unseen individuals, with the ability to personalize, (3) generate physiologically plausible heartbeat trajectories through forward simulation, and (4) remain physiologically interpretable to clinicians.

In this work, we address this gap by developing a framework using density-based neural temporal point processes (NTPPs) that combines the statistical rigor of point processes, physiological grounding of integrate-and-fire models of heartbeat generation, and the computational scalability of learned embeddings in a neural network, allowing a single model pretrained on a cohort containing both normal and arrhythmic populations to be personalized to an individual. Using five publicly available datasets spanning healthy, arrhythmic, atrial-fibrillation, and ischemic populations along with a realworld ambulatory dataset, we evaluate this framework on four tasks:

- **Scaling**: Formulating empirical scaling laws to establish how cross-subject pretraining improves held-out model calibration as cohort size grows.
- **Zero-shot transfer and personalization**: Evaluating how well the pretrained model generalizes to a held-out ambulatory dataset, with and without test-time adaptation.
- **Forward-simulation**: Demonstrating that our model sustains predictive accuracy and generates trajectories with physiologically plausible variability over short horizons.
- **Interpretability**: Exploring whether the internal learned representations can explain known physiological patterns. Together, these capabilities demonstrate how combining physiological rigor with scalable AI provides a practical blueprint for personalized cardiac monitoring and real-time forward simulation in clinical settings.

## II. Methods and Materials

### A. Problem Setup and Notation

Let 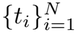denote heartbeat event times (R-peak times-tamps), with inter-event (RR) intervals *τ*_*i*_ = *t*_*i*_ −*t*_*i*−1_ *>* 0. A temporal point process over ordered events is characterized by the conditional density of the next waiting time, 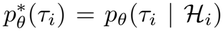, which is equivalent to the conditional intensity via *λ*^∗^(*τ*) = *p*^∗^(*τ*)*/*(1 − *F* ^∗^(*τ*)). We parameterize the density directly rather than the intensity, following [8], [9], which simplifies the negative log-likelihood. We minimize the negative log-likelihood to learn *θ* by:

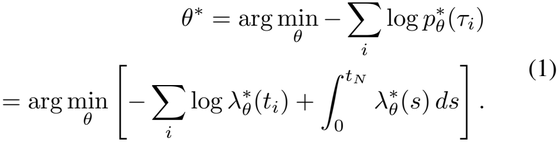

We parameterize 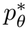as a mixture of log-normal components,

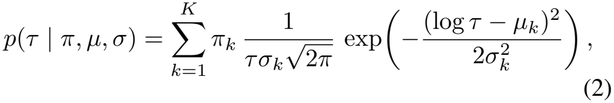

motivated by history-dependent inverse-Gaussian models established for RR intervals [1].

A gated recurrent unit encodes the history into a context vector, *c*_*i*_ = *h*_*i*−1_ with *h*_*i*_ = GRU(*x*_*i*_, *h*_*i*−1_), taking as input the standardized log-interval *x*_*i*_ = (log *τ*_*i*_− *μ*_*τ*_)*/σ*_*τ*_, where *μ*_*τ*_, *σ*_*τ*_ are computed over the training set. The mixture parameters are learned functions of the context (with *W, b* learned from data): *π* = softmax(*W*_*π*_*c*_*i*_+*b*_*π*_), *σ* = *σ*_*τ*_ exp(*W*_*σ*_*c*_*i*_+*b*_*σ*_), *μ* = *σ*_*τ*_ (*W*_*μ*_*c*_*i*_ + *b*_*μ*_) + *μ*_*τ*_, where the *σ*_*τ*_, *μ*_*τ*_ terms return the mixture from standardized log-time to the raw time scale. Since all three depend on *c*_*i*_, every parameter is history-dependent.

### B. Datasets

#### MIT-BIH Normal Sinus Rhythm Database (NSRDB)

Eighteen subjects (ages 20–50, 13 female), each providing ≈ 130 minutes of continuous ECG recording under normal sinus rhythm [10].

#### MIT-BIH Arrhythmia Database (MIT-BIH)

Forty-eight annotated records (from forty-seven subjects), forming a cohort comprising both normal and arrhythmic rhythms, providing a substantially more heterogeneous and clinically challenging population than the NSRDB.

#### Long-Term AF Database (LTAF)

Eighty-four long-term ECG records (≈24–25 hours each, two leads at 128 Hz) from patients exhibiting paroxysmal or sustained atrial fibrillation [13].

#### Long-Term ST Database (LTST)

Eighty-six continuous ECG records (≈ 21–24 hours each, two or three leads at 250 Hz) from eighty subjects, capturing transient ST-segment events and a broad range of ischemic and axis-shift episodes [14].

#### Icentia11k

541,794 ECG records (with ≈70 minute segments sampled from 3–14 days per patient, sampled at 250 Hz from a single lead heart monitor device) from 10,998 subjects, containing normal sinus, atrial fibrillation and atrial flutter rhythms [11].

#### Stanford 24-Hour Dataset

A dataset containing 10 ambulatory 24-hour ECG recordings collected under IRB approval from Stanford University via open-source non-invasive wearable hardware in real-world conditions. This dataset captures naturalistic physiological variation associated with daily activity, sleep, and meals, making it substantially harder than controlled short-duration recordings.

R-peak times are extracted using the Pan–Tompkins algorithm [15] for datasets that did not have manually annotated R-peaks.

### C. Evaluation Metrics

#### Goodness-of-fit testing

We use the Kolmogorov-Smirnov (KS) distance based on the Time Rescaling Theorem [9], [16] to measure distributional calibration. This measures whether predicted probability densities match true conditional distributions. If a model correctly captures the heartbeat generative process, the transformed inter-event intervals *z*_*i*_ should be independent and identically distributed uniform random variables on the interval (0, 1).

Let *τ*_*i*_ denote the *i*-th observed RR interval and H_*i*−1_ the history prior to event *i*. If *F* (·|H_*i*−1_) is the model’s conditional cumulative distribution function for the next RR interval, then the time-rescaled variables are

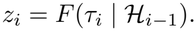

The KS-distance is defined as the maximum deviation between the empirical cumulative distribution of the *z*_*i*_ values and the CDF of a uniform distribution:

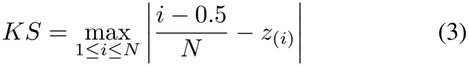

where *z*_(*i*)_ are the sorted transformed intervals. A well-fit model yields a KS-distance that falls within the 95% significance cutoff, defined as:

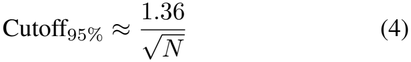

We define the % goodness-of-fit as the percentage of non-overlapping 2-minute epochs for which the KS-distance is within this cutoff.

#### Mean Absolute Error

For subject *s* at beat index *j* within a window, let 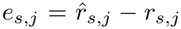denote the signed prediction error in seconds, where 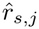and *r*_*s,j*_ are the predicted and actual RR intervals respectively.

To characterize point-wise error in a given horizon, we compute cumulative MAE curves. At beat index *k*, errors are averaged first within each subject and then across all *N* subjects:

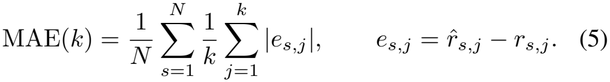

#### Root Mean Squared Error

RMSE is computed similarly, using

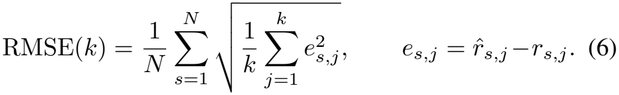

#### Traditional HRV measures

The SDNN (standard deviation of RR) measures the overall variability of the RR intervals, reflecting both short- and long-term components of heart rate variability. For RR intervals {*τ*_1_, …, *τ*_*N*_ } with mean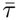:

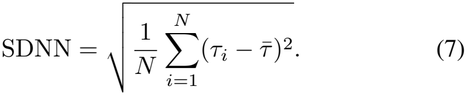

The RMSSD (root mean square of successive differences) quantifies short-term variability by measuring the root mean square of differences between successive RR intervals. With successive differences *d*_*i*_ = *τ*_*i*+1_ − *τ*_*i*_ (*i* = 1, …, *N* − 1):

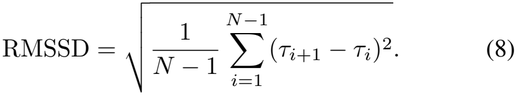

While KS distance and goodness-of-fit are the primary metrics used to evaluate our model, we additionally compute MAE and RMSE in our scaling and simulation experiments and RMSSD and SDNN in our simulation experiments.

### D. Formulating a Physiological Scaling Law

Neural scaling laws are empirical formulas that describe how model loss or error improves predictably as a powerlaw function when increasing data size, compute or model complexity. Establishing this allows us to quantitatively decide data collection budgets and determine optimal training budgets when training large models. To this end, we formulate a physiological scaling law describing how held-out metrics improve with the number of ECG recordings for pre-training, using a pooled corpus that combines healthy and arrhythmic cohorts (Icentia11k, NSRDB, MIT-BIH Arrhythmia, Long-Term AF, Long-Term ST). We ask (a) how held-out fit quality improves as pretraining subjects accumulate across the five datasets, and (b) whether a model trained on the pooled corpus generalizes evenly, which we assess by reporting goodness-of-fit (GOF), KS distance, RMSE, and MAE.

#### Pretraining scaling protocol

We pool 11,226 subjects from five datasets and representatively split them at the subject level into 10,184 training, 521 validation and 521 test subjects. Training subjects are added round-robin across the five datasets. Walking a log-spaced grid *n*∈ {1, 2, 4, …, 4096, 10184 }, a fresh NTPP is trained from scratch on the first *n* subjects using 2-minute streaming windows with up to 2 hours (60 windows) per subject drawn evenly across each recording (with the exception of MIT-BIH Arrhythmia which used its entire 30 minute recording). Each trained model is evaluated chunkwise without online updates on the held-out test subjects: over successive 2-minute chunks we compute the Kolmogorov–Smirnov (KS) distance between predicted and empirical inter-event distributions, the percentage of good-fit chunks (KS distance below the 5% significance cutoff 1.36*/* 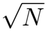), and one-step-ahead beat-wise errors (MAE, RMSE). We repeat this across 10 random seeds and report mean and standard deviation for all metrics.

Following [17] and our prior work [12], we fit each loweris-better metric (KS, MAE, RMSE) as a power law with an irreducible floor,

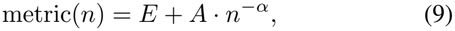

where *n* is the number of pretraining subjects, *E* is the irreducible floor (theoretic minimum error even with infinite data), *A* is the magnitude of initial error due to data scarcity, and *α* is the data-efficiency exponent that measures how performance improves as data size increases. For goodness-of-fit, where higher is better, we fit the same law to the gap 100− GOF, so its fitted ceiling is 100− *E*. We define **metric saturation** as the point where the reducible error component *A n*^−*α*^ falls within 1% of the irreducible floor *E*. Each (seed, grid) pair is one observation and we exclude *n* = 1 and *n* = 2, where single-subject effects dominate the fit, giving 12 grid points for each of the 10 seeds. Parameters are fit in log space, and are estimated using L-BFGS-B optimization with a robust Huber loss, which protects against fitting distortion from noisy empirical seeds at smaller *n* values. To quantify parameter uncertainty we use a seed-level (cluster) bootstrap (*samples* = 200).

### E. Test-Time Personalization via Online Adaptation

The base NTPP model can be used on its own, or serve as a pre-trained backbone that is adapted at test time to an individual subject to personalize predictions. We do this via online learning, updating the model weights for a brief period before testing, updating either all weights (*full model update*) or just the output layer weights (*head-only update*). We also test skipping the population baseline entirely and personalizing to the subject’s data alone. This gives four training paradigms:

- **V1: Population Baseline**. NTPP is trained offline on the five-dataset training cohort. Once trained, the model’s parameters are frozen and evaluated window-by-window on the test subject with no further updates. V1 isolates how well a static, population-level model generalizes to an unseen individual.
- **V2: Standalone Online Learning**. No pretrained model is used. We train a model initialized from scratch, calibrated on a brief burn-in segment of the test subject, which then learns online from its own stream. V2 tests whether a subject-specific model can be learned purely from streaming data, with no prior population knowledge.
- **V3.1: Population Pretraining + Full Online Adaptation**. The model is initialized from the V1 population base and then adapts online on the target subject’s stream, updating all parameters. After 25-minutes of adaptation, the adapted model is frozen and tested on streamed chunks. V3.1 tests whether combining population priors with individual adaptation beats either alone.
- **V3.2: Population Pretraining + Output-Only Online Adaptation**. Identical to V3.1, except online updates are restricted to the output layer only. This lighter-weight variant cuts per-update cost and reduces the risk of overwriting the pretrained representation (catastrophic forgetting).

We test the four paradigms on the held-out ambulatory Stanford dataset (10 subjects, our study uses 4 hour subsets for each recording). All updates use Adam with learning rate of 5× 10^−5^ with gradient clipping. We examine how online adapting the NTPP model affects distributional calibration metrics (KS distance and goodness-of-fit). All four paradigms are evaluated on the same test window. We report the mean ± 1 SD across subjects of chunk-wise KS distance and cumulative goodness-of-fit.

### F. Short-Horizon Autoregressive Forward-Simulation

We evaluate short-horizon autoregressive forecast performance of our model, where at test time, the model conditions only on its own generated outputs to predict future RR intervals. We compare both the static pre-trained model as well as output-only online adaptation for this experiment, comparing its trajectory to the true RR series, and evaluating by distributional calibration, point accuracy and variability. The frozen pre-trained model was trained on five-datasets (Icentia, NSRDB, MIT BIH, LTAF and LTST), with Icentia subjects dominating the training pool due to larger number of available training subjects. We perform training, validation and testing over 10 seeds and report the average ±1 SD for all metrics.

### G. Interpretability of the hidden state embedding and the lognormal mixture parameters

The GRU encodes the sequence of RR intervals into a 64-dimension hidden state, updated at each beat. We take the mean hidden state over non-overlapping 2-minute windows as that window’s local history vector. We explore whether the learned vectors naturally occupy different regions of the embedding space depending on the window’s broad rhythm annotation (normal or AFib/AFL) in the training set. We also ask whether these regions hold for unseen test windows, which can help in labeling a new RR sequence for a patient that the model has not seen prior. To transform the high-dimensional history vector into a 2-D visualizable form, we use t-SNE (t-distributed stochastic neighbor embedding), which maps distances between windows in the 64-dimensional space into neighbor probabilities and allows us to visualize emerging clusters based on t-SNE map coloring. We used openTSNE transform function to embed new out-of-sample data points onto the existing t-SNE embedding. The training subjects contribute far more Normal windows than AFib/AFL windows. The map therefore uses every AFib/AFL training window plus a random 50% of Normal training windows.

Each window was assigned one of two classes: Normal or Arrhythmia using rhythm annotations present in the dataset (for MIT BIH Arrhythmia, LTAF and Icentia11k) and we assume normal sinus rhythm for the remaining two datasets (NSRDB which is normal sinus rhythm and LTST which contains only ischemia or elevated ST segments). In order to test whether our test windows were labeled correctly, we visually inspect the embedded t-SNE map, but also quantify this using k-nearest-neighbor (kNN) classification in t-SNE space. Each embedded test window was assigned the majority class among its k=25 nearest training-map windows, weighted by the inverse of its class size.

We also analyze the NTPP model’s output distribution of the lognormal mixture parameters, to see if there is a statistically significant difference between the normal sinus and arrhythmia cohorts. For each input RR interval, the NTPP model outputs a mixture of K = 8 log-normal components with weights *w*_*k*_, location *μ*_*k*_ and scale *σ*_*k*_. For each beat we computed the mean and standard deviation of the predicted next-RR mixture. From these we derived the equivalent single log-normal mean and standard deviation (*μ*_*eff*_, *σ*_*eff*_). The spread and difference in the values were compared between the two groups using a two-sided Mann–Whitney U test. p-values were Holm-corrected within each family of related tests: mean, SD of the predicted RR, *μ*_*eff*_, *σ*_*eff*_ . Significance was set at *α* = 0.05.

## III. Results

### A. Pretraining Scales with Cohort Size

By fitting a scaling law to our pre-training scaling experiments as described in II-D, we show that model quality follows a power law with an irreducible floor (as seen in figure 1). The scaling-law coefficients are given in Table I. The fit for all four metrics are well-constrained(*R*^2^ = 0.94–0.97). Based on our saturation criteria, the fitted laws place saturation at roughly 1,100 subjects for RMSE, 2,400 for MAE, 3,200 for goodness-of-fit, and 3,700 subjects for KS distance. Extrapolating beyond our cohort to 100,000 subjects (Table II), KS distance is projected to fall from 0.0747 to 0.0744, goodness-of-fit rises from 93.0% to 93.3%, and MAE and RMSE each improve by less than 0.2 ms, showing no significant further gain.

**Fig. 1:**
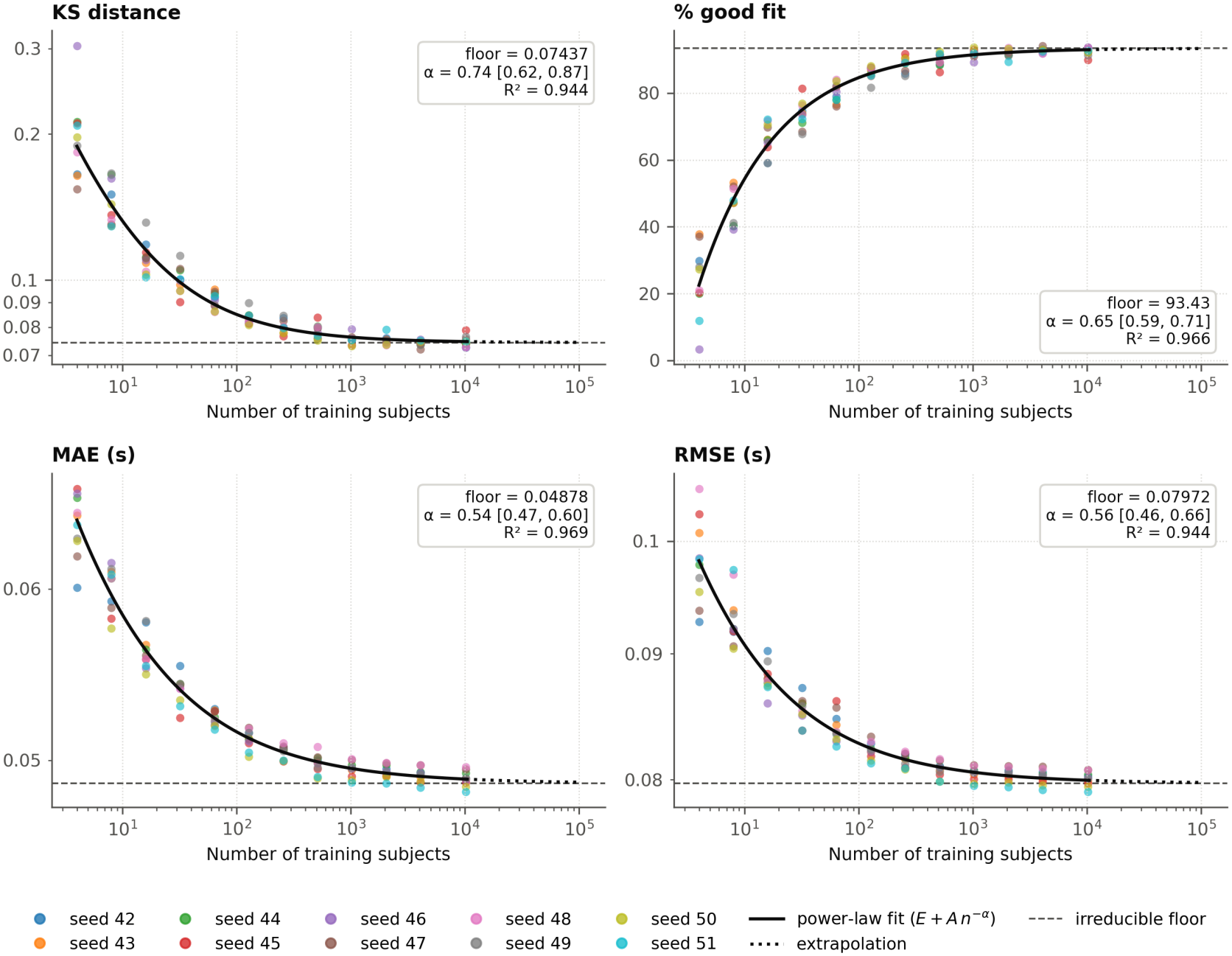
Held-out KS distance, %good-fit, beat-wise RMSE and MAE versus number of pretraining subjects (log–log; %good-fit on a linear *y*-axis). Points are 10 seeds × 12 subset sizes; solid curves are the fitted laws metric(*n*) = *E* + *A n*^−*α*^ and dashed lines the fitted floors *E*. We see that all four metrics follow power law distribution with an irreducible error component.

**TABLE I:**
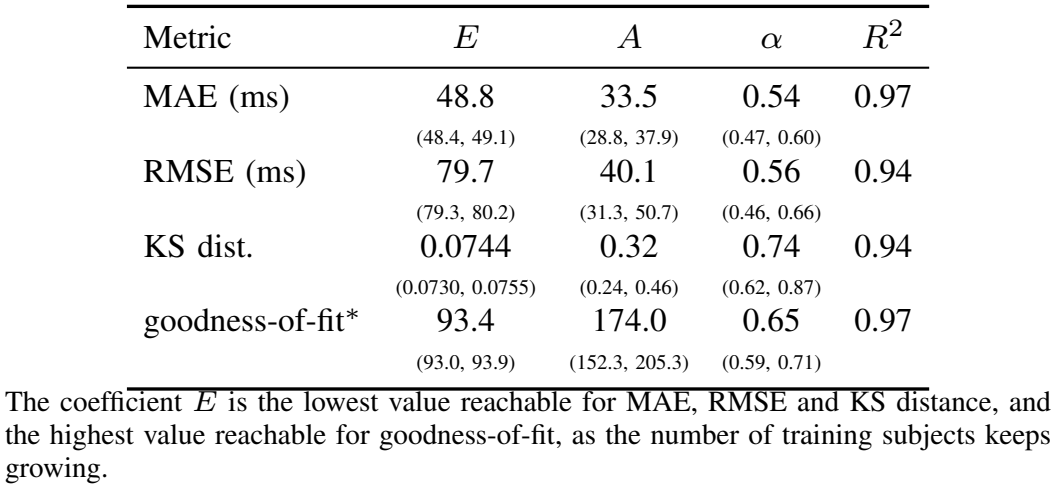
Fitted physiological scaling-law coefficients, *y*(*n*) = *E* + *A n*^−*α*^, where *n* is the number of training subjects. Values in parentheses are 95% CIs from 200 seed-level bootstrap resamples.

**TABLE II:** Projections from the fitted laws.

| Subjects *n* | KS dist. | goodness-of-fit | MAE (ms) | RMSE (ms) |
|---|---|---|---|---|
| 500 | 0.0776 | 90.3 | 49.93 | 80.96 |
| 1,000 | 0.0763 | 91.4 | 49.57 | 80.56 |
| **10,184** | 0.0747 | 93.0 | 49.00 | 79.95 |
| 50,000 | 0.0745 | 93.3 | 48.87 | 79.82 |
| 100,000 | 0.0744 | 93.3 | 48.84 | 79.79 |
| *n* → ∞ | 0.0744 | 93.4 | 48.78 | 79.72 |

### B. Population pre-trained model transfers zero-shot to held out dataset, and personalization gives marginal gains

Fig. 2 shows chunk-wise KS distance and cumulative goodness-of-fit over the Stanford test dataset. Table III summarizes quantitative results.

**Fig. 2:**
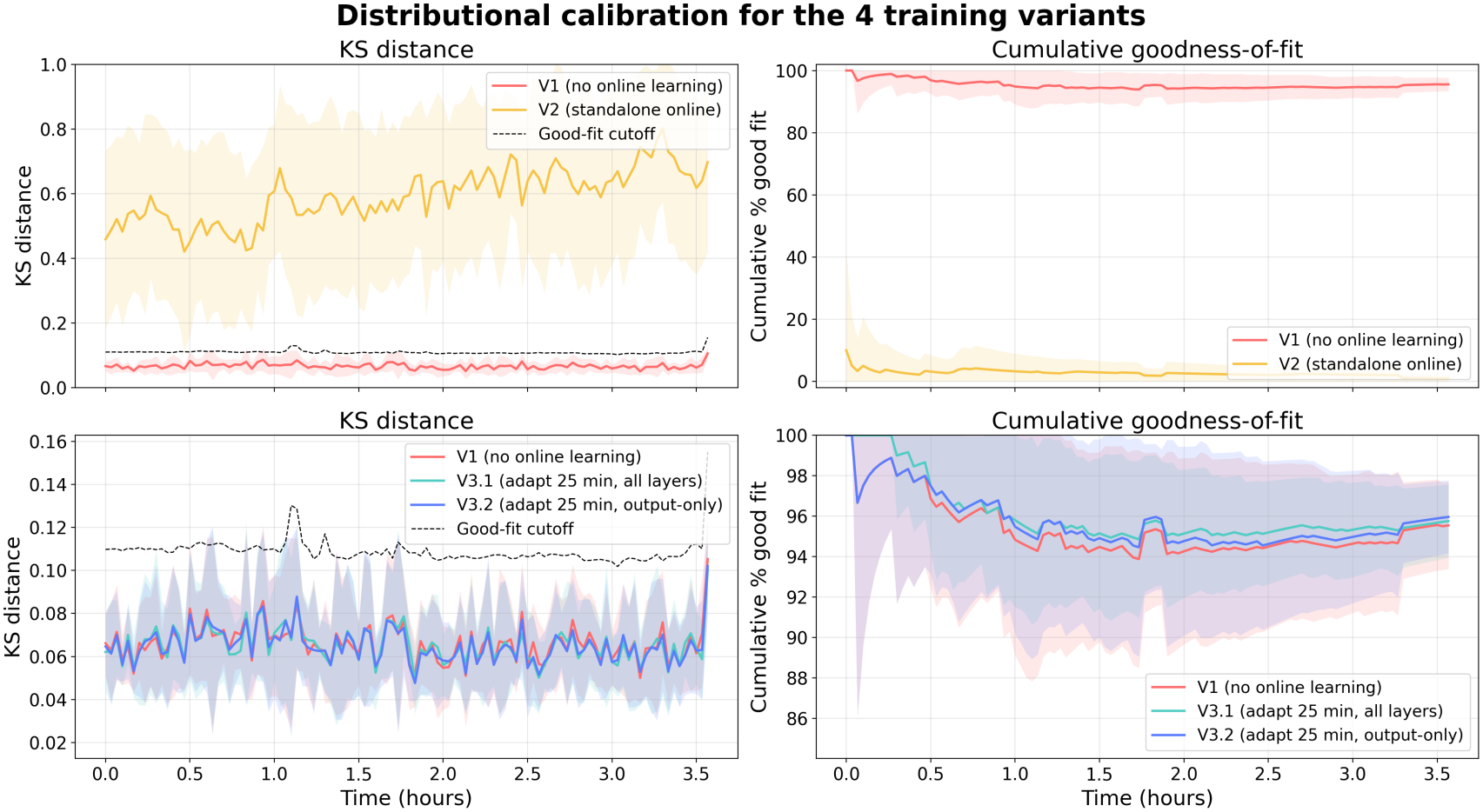
Zero-shot vs standalone training vs personalization on the realworld ambulatory Stanford dataset. Standalone online training on test subjects data fails compared to zero-shot pre-trained model. Adding test subjects data via 25-minutes of online adaptation only marginally improves metrics, showing that the pre-trained data on its own provides a strong baseline, even for unseen real-world ambulatory recordings. *(Top)* V1 vs V2. *(Bottom)* V1 vs V3.1 and V3.2.

**TABLE III:**
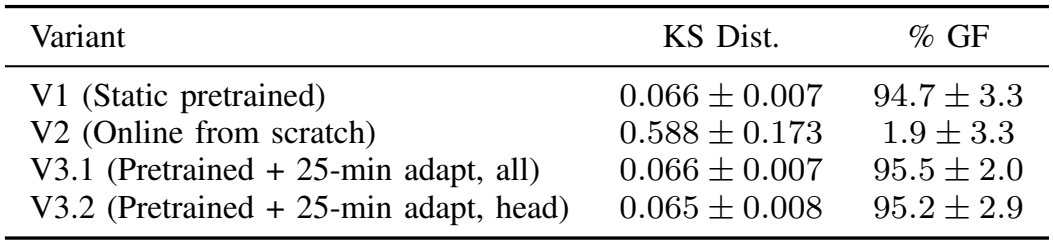
Online adaptation results on the held-out Stanford 4-hour recordings. Mean ± SD over *N* = 10 subjects.

| Variant | KS Dist. | % GF |
|---|---|---|
| V1 (Static pretrained) | 0.066 ± 0.007 | 94.7 ± 3.3 |
| V2 (Online from scratch) | 0.588 ± 0.173 | 1.9 ± 3.3 |
| V3.1 (Pretrained + 25-min adapt, all) | 0.066 ± 0.007 | 95.5 ± 2.0 |
| V3.2 (Pretrained + 25-min adapt, head) | 0.065 ± 0.008 | 95.2 ± 2.9 |

Pretraining on over 10,000 subjects gives us a model that is already well calibrated, even for subjects it has not seen. We see that online adaptation brings little to no benefit over the population baseline. V1 attains 94.7% goodness-of-fit (KS 0.066), and the adapted variants V3.1 and V3.2 reach 95.5% and 95.2% (KS 0.066 and 0.065). In contrast, online learning without pretraining (V2) collapses, reaching only 1.9% goodness-of-fit with a KS distance of 0.59 showing that a shared representation learned from a cohort of subjects is necessary.

Head-only adaptation (V3.2) updates only ∼ 11% of the network’s parameters (1,560 of 14,488) yet performs on par with full-model adaptation (V3.1), with near-equal goodness-of-fit and KS distance. When adaptation is used, head-only updating remains preferable: it needs less memory and compute, and limits the risk of overwriting the pretrained representation (catastrophic forgetting).

### C. Pretrained model remains calibrated over short forecast horizons

Figure 3 shows that point errors RMSE and MAE are relatively low and gradually increase across the horizon. Looking at distributional calibration(figure 4), we see that KS distance stays within the cutoffs (calculated using 1-minute windows) and maintains an average 87-88% goodness-of-fit over five minutes, and ends with a goodness-of-fit of 78-80% at the end of the trajectory. This shows overall the trajectory is well-calibrated, given that the model conditions on its own generated beats.

**Fig. 3:**
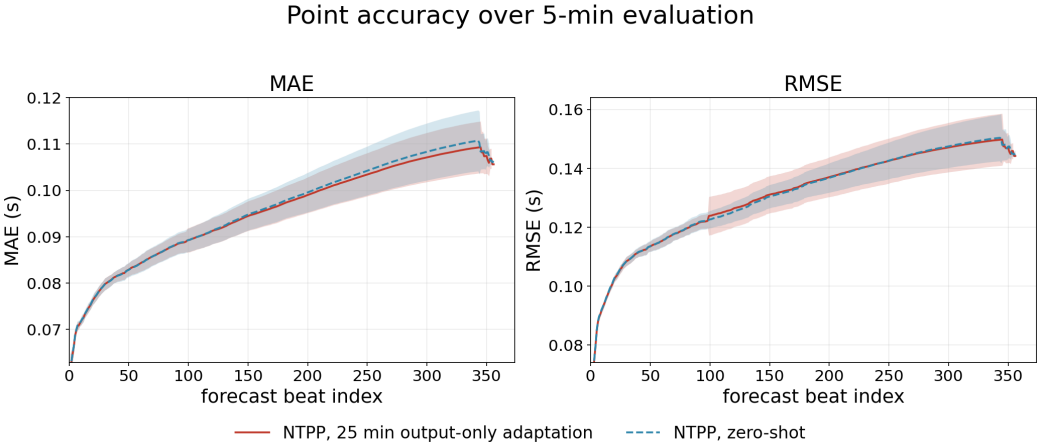
Beat-wise cumulative error MAE (left) and RMSE (right) show relatively low overall error, with gradual increase over the horizon. Red line corresponds to metrics following 25 min online adaptation, and blue dashed line corresponds to zero-shot metrics.

**Fig. 4:**
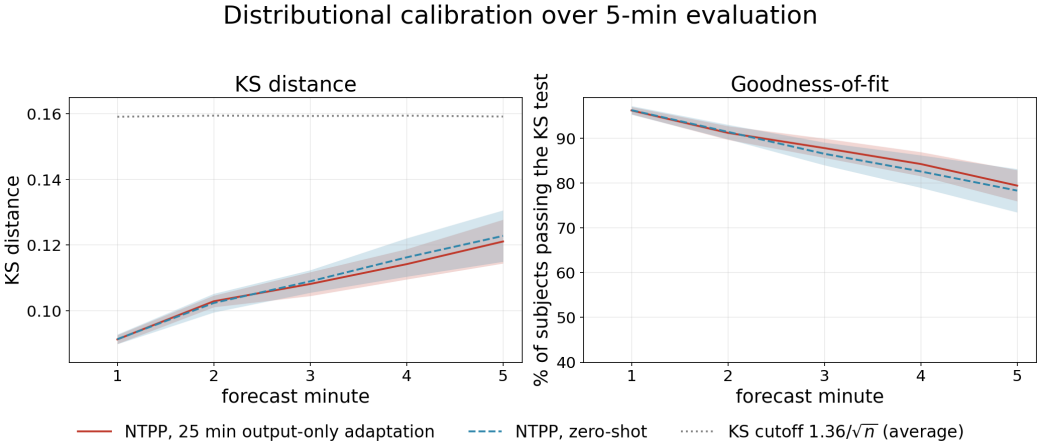
Minute-wise KS distance (left) and goodness-of-fit (right) averaged over test subjects across 10 seeds. Red line corresponds to metrics following 25 min online adaptation, and blue dashed line corresponds to zero-shot metrics.

Another important aspect of this experiment is to evaluate if the low absolute errors come from a realistic forecasted trajectory. Measuring the forecast variability is necessary, as we can get low errors by just predicting the mean, but this would not be meaningful, as seen in previous experiments applying large time series foundation models to RR interval forecasting [7]. Our experiments show that local variability (measured by SDNN and RMSSD with 30-beat rolling windows) is retained in our trajectory as seen in Figure 5. However, the variability in our forecasted trajectory is notably higher than that of the true RR series, which is a limitation.

**Fig. 5:**
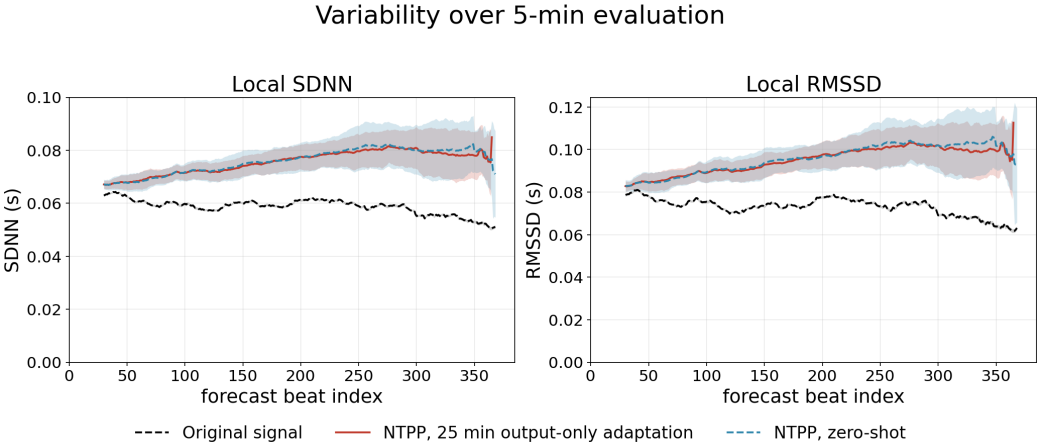
Beat-wise short-term heart-rate variability over the 5-minute forecast horizon, computed with a 30-beat rolling window with SDNN (left) and RMSSD (right). Black line corresponds to original RR series, red line corresponds to metrics following 25 min online adaptation, and blue dashed line corresponds to zero-shot metrics.

In general, we see that there are no significant differences in the errors between the zero-shot and online-adapted models in all evaluation metrics. A summary of the performance metrics by dataset is given in Appendix Table IV.

**TABLE IV:** Forward simulation performance over the 5 minutes (mean ± SD across the 10 seed models), using zero-shot and output-only online adaptation on 25 min of data.

| **Dataset** | **Model** | **MAE (s)** | **RMSE (s)** | **KS distance** | **Goodness of fit** |
|---|---|---|---|---|---|
| NSRDB | 25-min adaptation | 0.140 ± 0.077 | 0.173 ± 0.083 | 0.117 ± 0.030 | 84% |
|  | Zero-shot | 0.146 ± 0.078 | 0.180 ± 0.085 | 0.120 ± 0.034 | 79% |
| LTAF | 25-min adaptation | 0.141 ± 0.027 | 0.187 ± 0.034 | 0.118 ± 0.015 | 80% |
|  | Zero-shot | 0.148 ± 0.032 | 0.194 ± 0.040 | 0.119 ± 0.012 | 78% |
| LTST | 25-min adaptation | 0.101 ± 0.024 | 0.136 ± 0.027 | 0.113 ± 0.022 | 82% |
|  | Zero-shot | 0.103 ± 0.025 | 0.137 ± 0.030 | 0.112 ± 0.019 | 83% |
| Icentia11k | 25-min adaptation | 0.111 ± 0.005 | 0.152 ± 0.009 | 0.107 ± 0.003 | 88% |
|  | Zero-shot | 0.112 ± 0.006 | 0.153 ± 0.008 | 0.108 ± 0.004 | 87% |
| Overall | 25-min adaptation | 0.111 ± 0.006 | 0.152 ± 0.009 | 0.108 ± 0.003 | 88% |
|  | Zero-shot | 0.113 ± 0.007 | 0.153 ± 0.009 | 0.108 ± 0.004 | 87% |

### D. Learned Representations Encode Rhythm Irregularity

Figure 6 shows that the training map splits into a normal region and an Arrhythmia region. New unseen test windows land in the region of their own category. We also use kNN on the t-SNE space to predict the class label for unseen test windows, which gives us a balanced accuracy of 95.7% (with *recall*_*Normal*_ = 0.988; *recall*_*Arrhythmia*_ = 0.927).

**Fig. 6:**
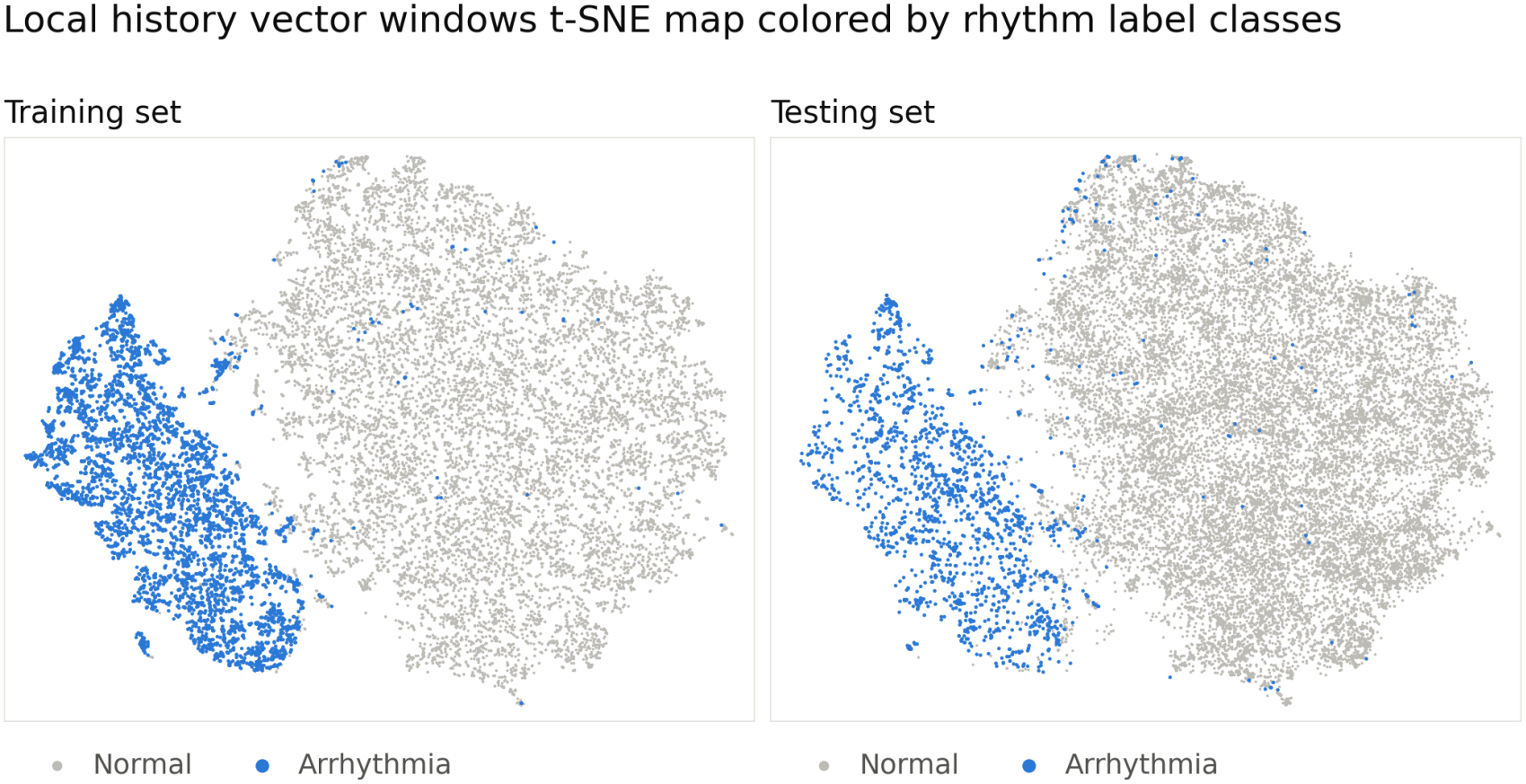
Local history vectors of 2-minute RR sequences, mapped with t-SNE and colored by rhythm label class. Left subplot is training data (Arrhythmia includes all points, 50% of Normal windows were sampled) and right subplot is the test windows embedded onto the map.

Figure 7 shows that the mean of the next predicted distribution (mean RR) and the mean of the single log-normal (*μ*_*eff*_) show lower values in the Arrhythmia class than normal, and also show large difference in their spread (standard deviation of the next predicted distribution and single log-normal).

**Fig. 7:**
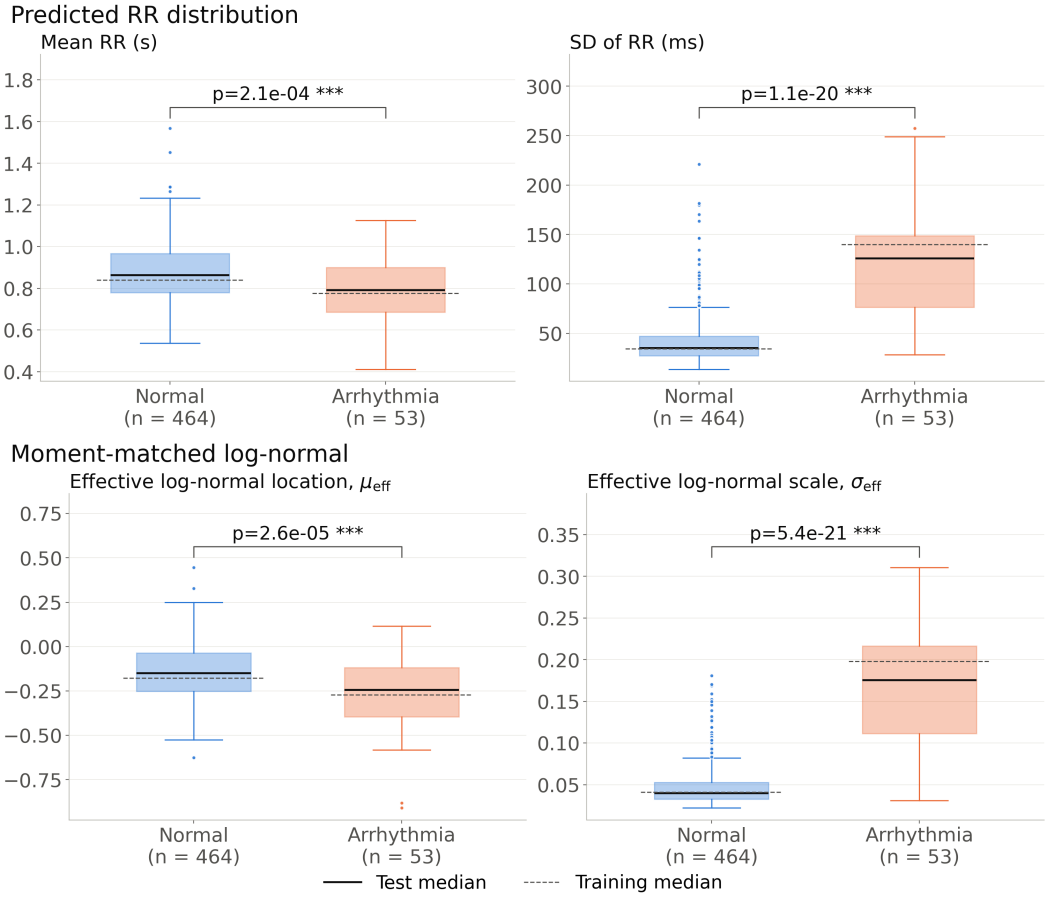
Top row: mean and standard deviation of the model’s predicted next-RR mixture. Bottom row: location (*μ*_*eff*_) and scale (*σ*_*eff*_) of the single log-normal with the same mean and SD as the mixture. Boxes show the interquartile range (IQR), solid lines show held-out medians; dashed lines show training medians.

The arrhythmia cohort had a slightly shorter, but statistically significant, predicted RR interval than the normal group (median 0.79 vs. 0.86 s; *p* = 2.1 ×10^−4^). The effective location *μ*_eff_ of the lognormal mixture follows the same pattern, with lower values in the arrhythmia group (*p* = 2.6 × 10^−5^). The standard deviation of the predicted next-RR distribution was about 3.6× larger in arrhythmia than in normal (126 vs. 35 ms; *p* = 1.1×10^−20^), and the effective spread *σ*_eff_ was about 4.4× larger (0.175 vs. 0.040; *p* = 5.4 × 10^−21^).

## IV. Discussion

In this study, we present a probabilistic foundation model for heartbeat dynamics from single-lead ECG by training a density-based NTPP on RR-interval sequences from over 11,000 subjects. There are four main takeaways from our work.

First, our scaling experiments show that held-out metrics improve with increase in cohort-size, following a power law with irreducible floor. Extending our earlier 183-subject analysis [12] to over 10,000 subjects, we confirm that scaling in stochastic-output domains should be judged by distributional calibration. Point errors have a large irreducible component and change little across cohort sizes (MAE fell by ≈ 24%), whereas KS distance fell by ≈ 60% and goodness-of-fit rose from 25% to 93%.

Second, our population pre-trained model is well calibrated on an unseen ambulatory cohort, with 94.7% goodness-of-fit. Test-time adaptation of all weights or only the output layer gave marginal gains, and a newly initialized model trained only on the test subject’s own data fails catastrophically (1.9%), showing that a base pretrained model is necessary.

Third, in autoregressive forward simulation over short horizons, our model sustains predictive accuracy and calibration, while capturing local heart rate variability. We show that our model preserves the physiologically plausible beat-to-beat dynamics, although our variability is higher than the true series.

Lastly, in our interpretability experiments, we show that the learned representations, both the hidden state vector and the parameters of the lognormal mixture, are physiologically interpretable. The GRU history embedding organizes 2-minute windows by rhythm class without the model having any knowledge of rhythm or beat annotations, and this generalizes to held-out test subjects (95.7% balanced accuracy). In the output lognormal distribution, arrhythmic windows differ from normal windows mainly in predictive spread (roughly 3.6x larger predicted SD), with a much smaller shift in predicted mean RR.

There are some limitations to our study. First, due to compute constraints, our scaling experiments were limited to using only 2-hour recordings per subject rather than full-length traces. Second, our current forward-simulation au-toregressively rolls out beats, sampling from the lognormal distribution, but this leads to higher variability. Finally, our current framework focuses exclusively on single-lead ECG inter-beat intervals, leaving multimodal signals unexplored.

By combining the biological principles of cardiac pacemaking with the capabilities of modern AI, this work offers a physiologically grounded and scalable framework for continuous monitoring. This lays the groundwork towards providing clinicians with better predictive insights and supporting the development of digital twins and patient simulations for clinical research.

## V. Appendix

### A. Forward simulation evaluation metrics

